# Categorizing SHR and WKY rats by chi2 algorithm and decision tree

**DOI:** 10.1101/2020.02.03.931899

**Authors:** Ping-Rui Tsai, Kun-Huang Chen, Tzay-Ming Hong, Fu-Nien Wang, Teng-Yi Huang

## Abstract

In the past two decades neuroscience has offered many popular methods for the analysis of mental disorder, such as seed-based analysis, ICA, and graph methods. They are widely used in the study of brain network. We offer a new procedure that can simplify the analysis and has a high ROC index over 0.9. This method uses the graph theory to build a connectivity network, which is characterized by degrees and measures the number of effective links for each voxel. When the degree is ranked from low to high, the network equation can be fit by the power-law distribution. It has been proposed that distinct and yet robust exponents of the power law can differentiate human behavior. Using the mentally disordered SHR and WKY rats as samples, we employ chi2 algorithm and Decision Tree to classify different states of mental disorder by analyzing different traits in degree of connectivity.

## Introduction

Recent studies showed that human brain actions can be expressed by different network equations^1^ with the aid of graph method by fMRI samples^2^. Power law has been reported for many complex physical systems. Examples are the city population^3^, world wide web^4^, fluctuations in financial market^5^ et al. In this paper, we discuss the power law trait of authorized^6,7,8^ rat samples with SHR and WKY. Isoflurane is used to further divide our samples into four states: high isoflurane WKY=HW, high isoflurane SHR=HS, low isoflurane SHR=LS, and low isoflurane WKY=LW. The format of our sample is 11 slices, 525 times, 64×64, FOV=30mm, and slice thickness=1 mm. And each state contains 20 data.

We use the same procedure as ref.1 to obtain the power-law distribution for the fMRI data of rats. Pearson correlation defined in Eq.1 plays an important role because this calculable quantity can reflect the strength of positive correlation between any two voxels:

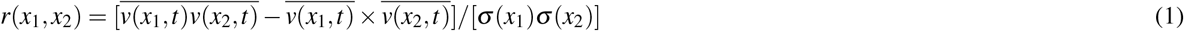

where a voxel at position *x* and time *t* is denoted as *v*(*x, t*) while *σ* represents the standard deviation. When Eq.1 exceeds a threshold value, chosen to be 0.7, these two voxels are regarded as being linked. After statistically analyzing all different degree of connectivity in the whole brain, we can obtain the power law distribution. Figure 1 shows the average distribution of these four states. Power law distribution is extremely useful because it sheds light on the difficult problem of analysing mental states. But it is insufficient to merely use its exponent to distinguish samples because ROC index is always less than 0.7. Based on this reason, chi2 algorithm will be quoted to help us select the significant difference from a group of degrees. After chi2 algorithm, C4.5 decision tree can help us produce a tree structure. Our purpose is to demonstrate whether C4.5 can offer a better way to help us observe and detect power law. Decision tree is a very popular tool for classification in data mining, which is widely used in deep-learning and machine learning^9,10^, industrial application^11,12^, medical treatments^13,14,15^, and bioinformatics^16,17,18^. To familiarize the readers with how decision tree can be of use in practical problems, let’s imagine if we want to know who was dead among the passengers who boarded the Titanic. First, we can quote the list of passenger, such as gender, age or level of class on the boat. Second, using this list to make decision tree. Finally, decision tree will tell us which condition can effect the fate of each passenger. Obviously, in this paper power law is the analogy of the list - different degrees are like the factors.

**Figure 1.**
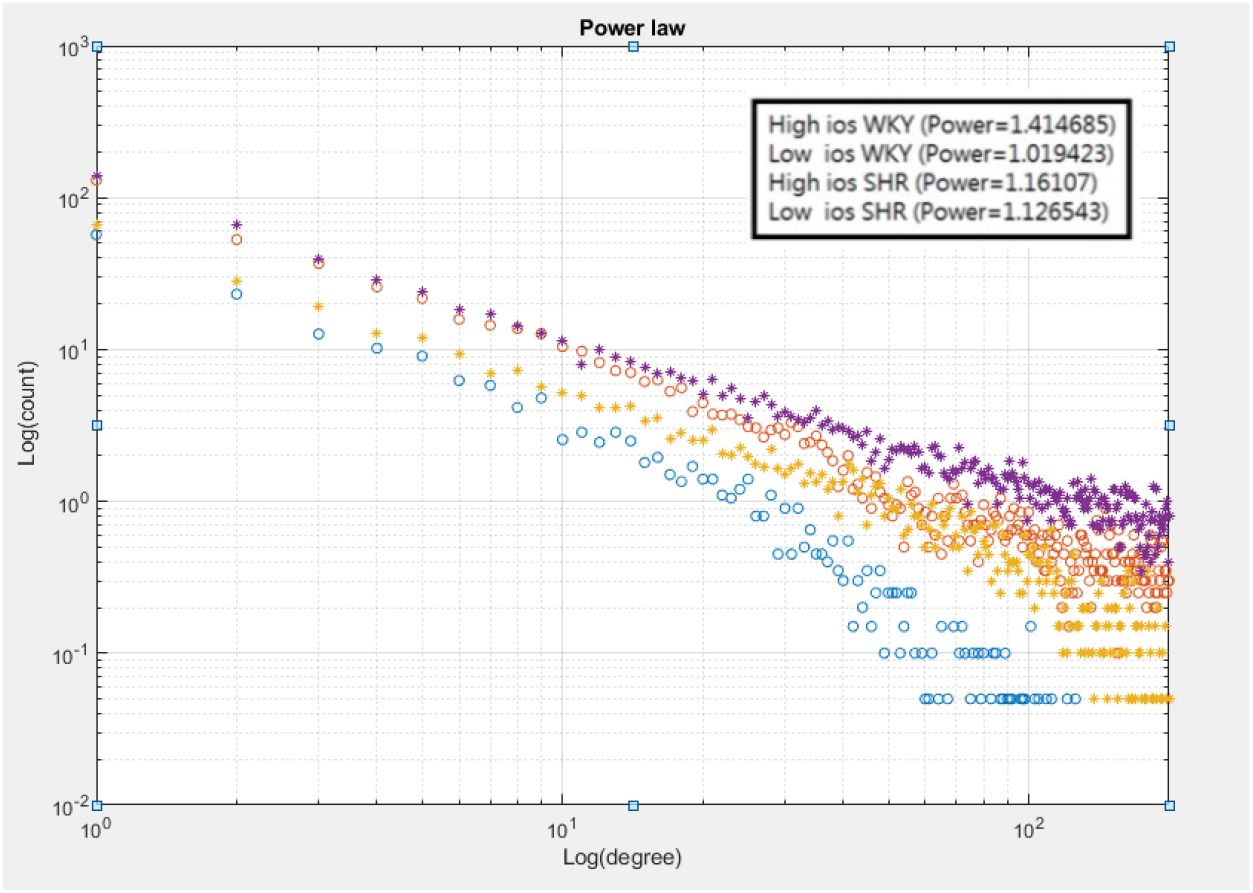
The count versus degrees of connectivity shows a power-law behavior. Each state has 20 samples. Hw and Hs are represented by blue and orange circles, while Lw and Ls are yellow and purple stars.

In the Result section of this paper, we will ensure the relation between C4.5 decision tree and the power law, such as a bar code and detector. This method can be a good starting point to establish a dynamical system to describe different mental states by evolution of degree. For processing, we use MATLAB to deal with fMRI raw data. The GPU and CPU mixing program allows us to increase the execution efficiency. The former transforms the 4D (3D voxel space and time) into a 2D matrix (voxel position and time), while the latter handles the calculation of correlation. As for the decision tree, we save the MATLAB matrix file by csv format, input the data into Excel, and output to Java to build the decision tree for the final calculation of the outcome of 10-fold cross-validation.

## Results

Table 1 shows the important features from chi2 algorithm, while Fig.2 shows the degree distribution after chi2 algorithm. By comparing with Fig.1, one can see that Fig.2 manages to widen the separation between degree distributions. Now we can input important features into C4.5 by 10-fold cross-validation before selecting the tree whose ROC approaches 0.9. Table 2 shows that three of our results in fact manage to obtain ROC > 0.9. Figure 3A explains how the four states are distinguished. Initially, data are categorized into two groups, based on whether the dosage of isoflurane(iso) ≥ 2.0. Figures 3B and 3C show results from different situations of high ios and low ios groups. Our data are processed by 10 fold cross-validation to achieve good results with ROC > 0.8. We double-confirm that power law has high sensitivity to symbolize rat problem. The result turns out to be positive which implies that power law is indeed a good representation for analysis.

**Figure 2.**
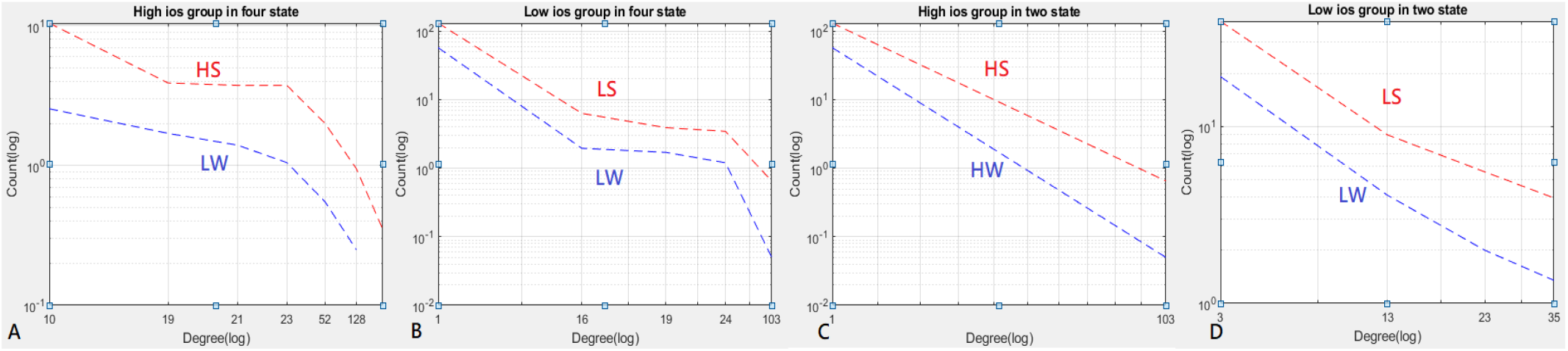
These figures show the power law distribution after chi2 algorithm.

**Figure 3.**
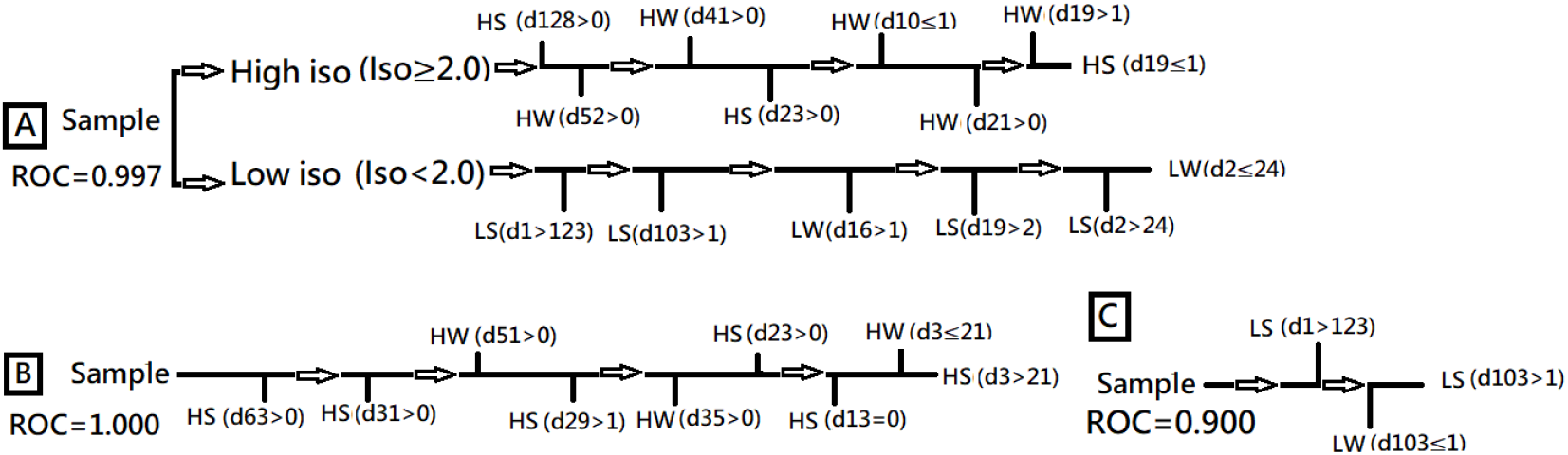
3A: Decision tree of HS, HW, LS, and LW states. 3B: Decision tree of HS and HW states. 3C: Decision tree of LS and LW states.

**Table 1.** Selected data by chi2 algorithm from the power law.

| Proposition | Degrees (after chi2 algorithm) |
| --- | --- |
| 4 groups (high/low iso WKY and SHR) | High iso: 128, 52, 41, 23, 10, 21, 19; Low iso: 1, 103, 16, 19, 24 |
| 2 groups (high iso WKY and SHR) | 1, 103 |
| 2 groups (low iso WKY and SHR) | 63, 31, 51, 29, 3, 35, 23, 13, 3 |

**Table 2.** ROC index for three different Decision Trees.

| Proposition | Degrees (after chi2 algorithm) | Number of samples |
| --- | --- | --- |
| 4 groups (high/low iso WKY and SHR) | 0.997 | 80 |
| 2 groups (high iso WKY and SHR) | 1.000 | 40 |
| 2 groups (low iso WKY and SHR) | 0.900 | 40 |

## Discussion

Figure 4 shows some representative properties of graph method of L which is defined as the length of links between any two nodes with the unit of mm and is important in small-world structure^19,20^. We found that, regardless of whether the sample is of high or low ios, the average and maximum L-values of SHR are always larger than those of WKY, as demonstrated by Fig.4A and 4B. The average nonzero degrees between 1 and 200 in Fig.4C and the link number in Fig.4D also appear to be greater than those of WKY. These observations may explain why SHR samples are more hyperactive than their WKY counterparts.

**Figure 4.**
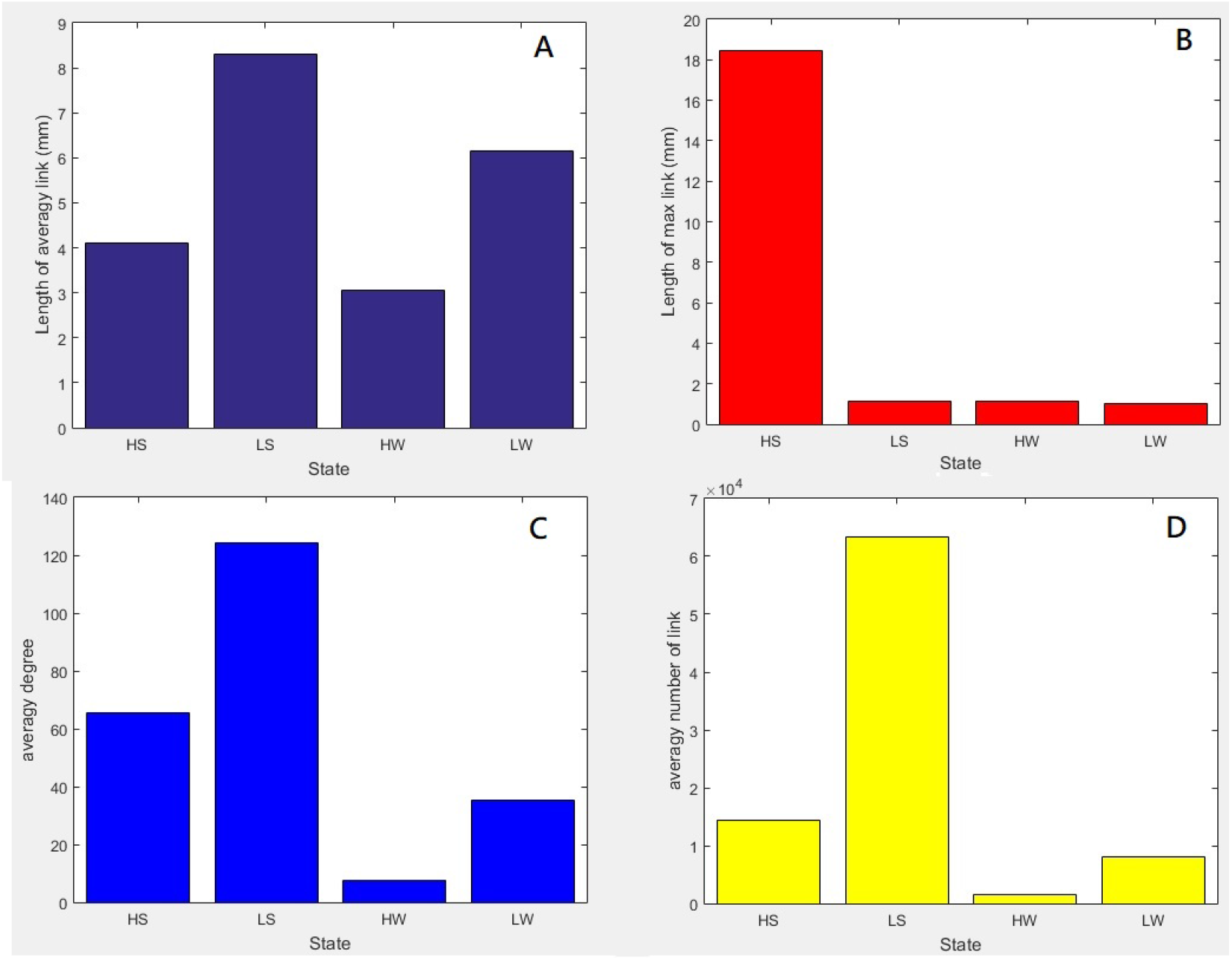
Important information of L between any two nodes that share a common connection. Panel A shows the average L, B the maximum L for each state, C the average degree, and D the average number of links. Each state has 6 samples.

Figure 5 shows that all states have different degree distribution. Only the most representative result is selected among 20 rat samples for each state. In general, SHR has more degrees and covers more brain regions than WKY. The degree distribution for low iso samples has more degrees and covers more regions than high iso. Table 3 shows the rank of activated brain region. Whenever the brain develops disorder, it exhibits a different functional network that consequently gives rise to a new exponent for the power law in Fig.1. This is similar to the finding of Ref.1 that the exponent may vary as the trial subject engages in different activities. We can find that the most active brain regions are the same. However, if we just focus on the samples of SHR, we discover that the secondary motor^21^ will fall to rank four. This is the reason why LS rat is more active than HS rat. More details will rely on more biological experiments in the future. The prefrontal cortex of ADHD patients has been reported to show abnormalities^22,23^. In our case, we can check two important regions in the prefrontal cortex to take states apart. These regions, Prl^24^ and Fra^25^, are related to the self-control and ADHD. The prefrontal cortex of SHR rat has been studied^26,27,28^. Figure 6 shows the number of degrees connecting other brain regions. We expect either Prl or Fra can give specific evidence expressing different characteristics for the four states in Table 4 by t-test. We found in Table 4 that Prl and Fra should be combined in order to give p-value that is less than 0.1. Figure 6 shows the average of degrees in Prl and Fra. Note that the stimulus interaction for LS being lower than LW in Fra is contrary to our expectation. Future biological experiments are needed to clarify the source of this problem.

**Figure 5.**
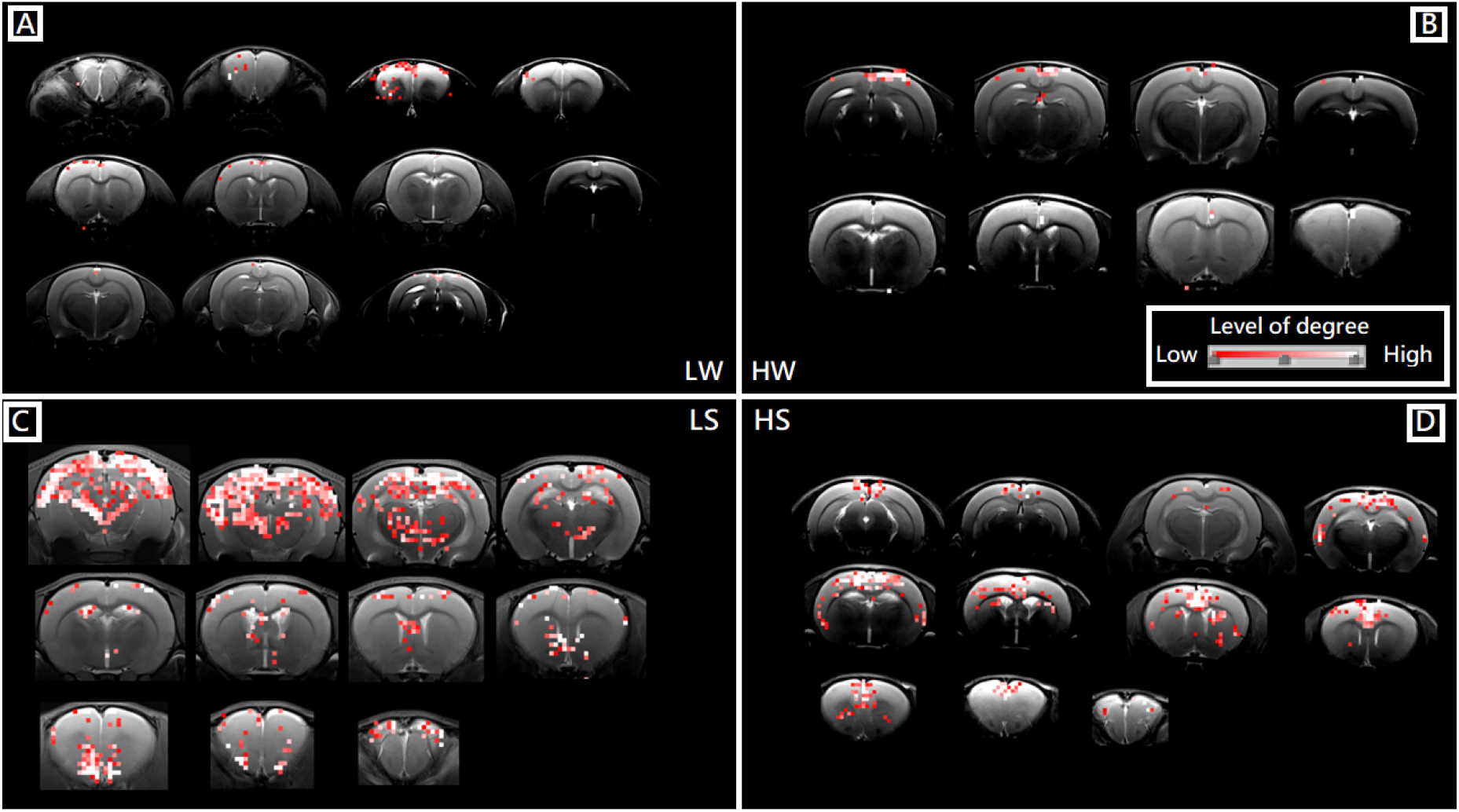
Panels A to D illustrate the activated region in LW, HW, LS, and HS states, respectively. The grade of brightness signifies different degrees of connectivity in the cross section. We just show the brain section with points.

**Figure 6.**
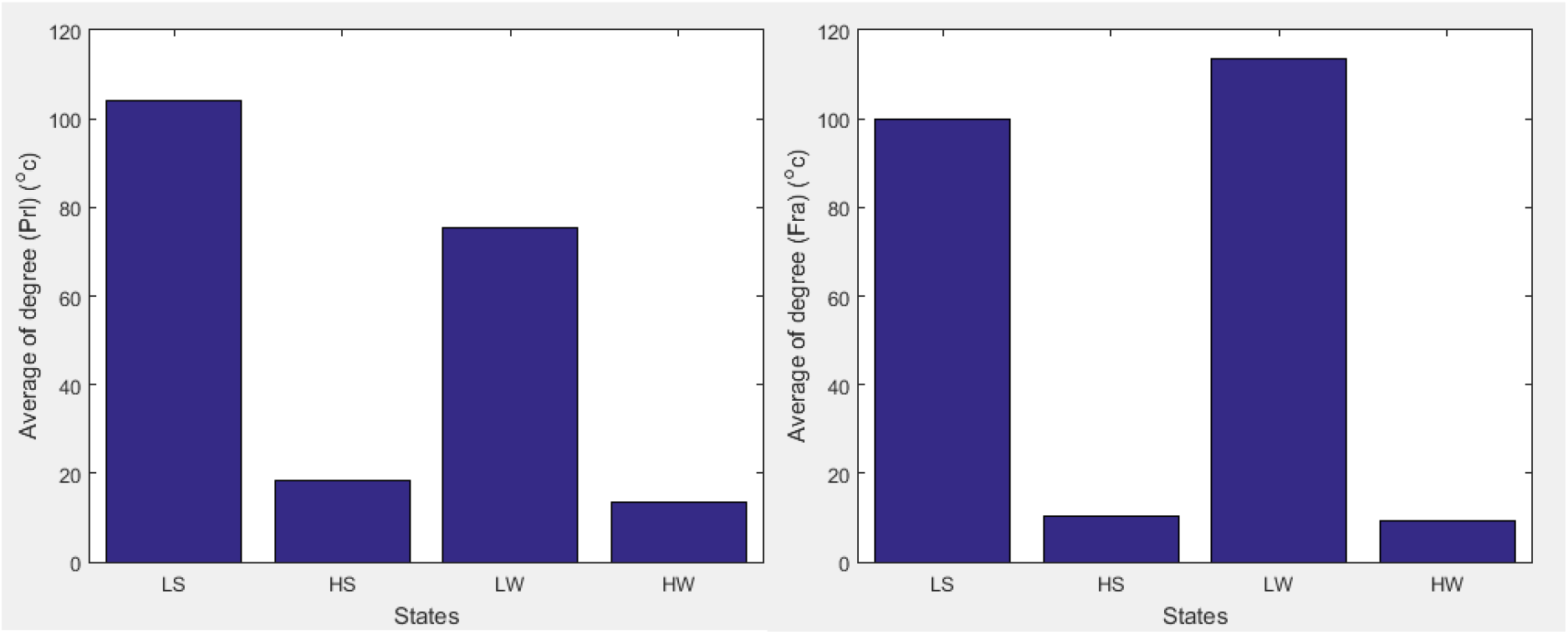
The average of degrees is calculated for Prl and Fra. Both figures give a higher average for low iso, as expected. But only Prl gives the right trend of a higher average for LS than LW.

**Table 3.** According to Fig.5, we list the name of brain areas that are activated. The brain area that exhibits the same activated frequency is noted in the parentheses.

| Rank | LW | HW |
| --- | --- | --- |
| 1 | Secondary motor cortex | Secondary motor cortex |
| 2 | Primary motor cortex | Primary motor cortex |
| 3 | Cingulate cortex, area 1 | Cingulate cortex, area 1 |
| 4 | Retrosplenial agranular cortex | Primary somatosensory cortex, barrel field |
| 5 | Retrosplenial granular b cortex | Primary somatosensory cortex, jaw region (rank 4) |
| Rank | LS | HS |
| 1 | Secondary motor cortex | Primary motor cortex |
| 2 | Primary motor cortex (rank 1) | Retrosplenial agranular cortex |
| 3 | Cingulate cortex, area 1 | Cingulate cortex, area 1 |
| 4 | Retrosplenial agranular cortex | Secondary motor cortex |
| 5 | Primary somatosensory cortex, dysgranular region | Retrosplenial granular b cortex (rank 4) |

**Table 4.** The p value for different states is calculated from Fra or/and Prl brain region.

| T-test | p-value of Fra | p-value of Prl | p-value of mixing Fra and Prl |
| --- | --- | --- | --- |
| HN - HS | 0.239 | 0.000 | 0.000 |
| HN - LW | 0.000 | 0.000 | 0.000 |
| HN - LS | 0.000 | 0.000 | 0.000 |
| HS - LW | 0.000 | 0.345 | 0.000 |
| HS - LS | 0.000 | 0.000 | 0.000 |
| LW - LS | 0.156 | 0.000 | 0.000 |

Clustering coefficient C is a measure to gauge how likely different nodes will form a cluster. This index is common for describing network structures, such as Fig.7. The definition for C is C=2N/[D(D-1)] where D denotes the degree for each voxel, while N is the maximum number of links among its neighbors. From the different clustering coefficients in Fig.8, we can conclude that the arrangement of nodes and edges are sensitive to the nature of mental disorder states.

**Figure 7.**
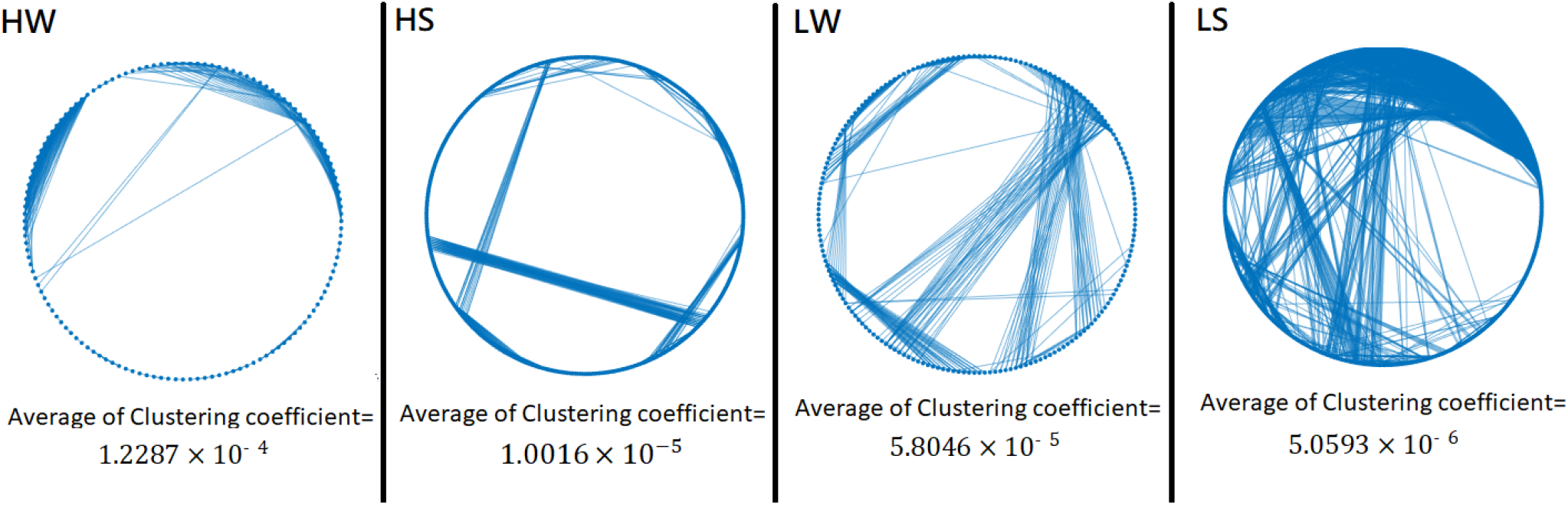
Comparing C and network structure for HS, LS, HW, and LW states. SHR has more functional connections than WKY.

**Figure 8.**
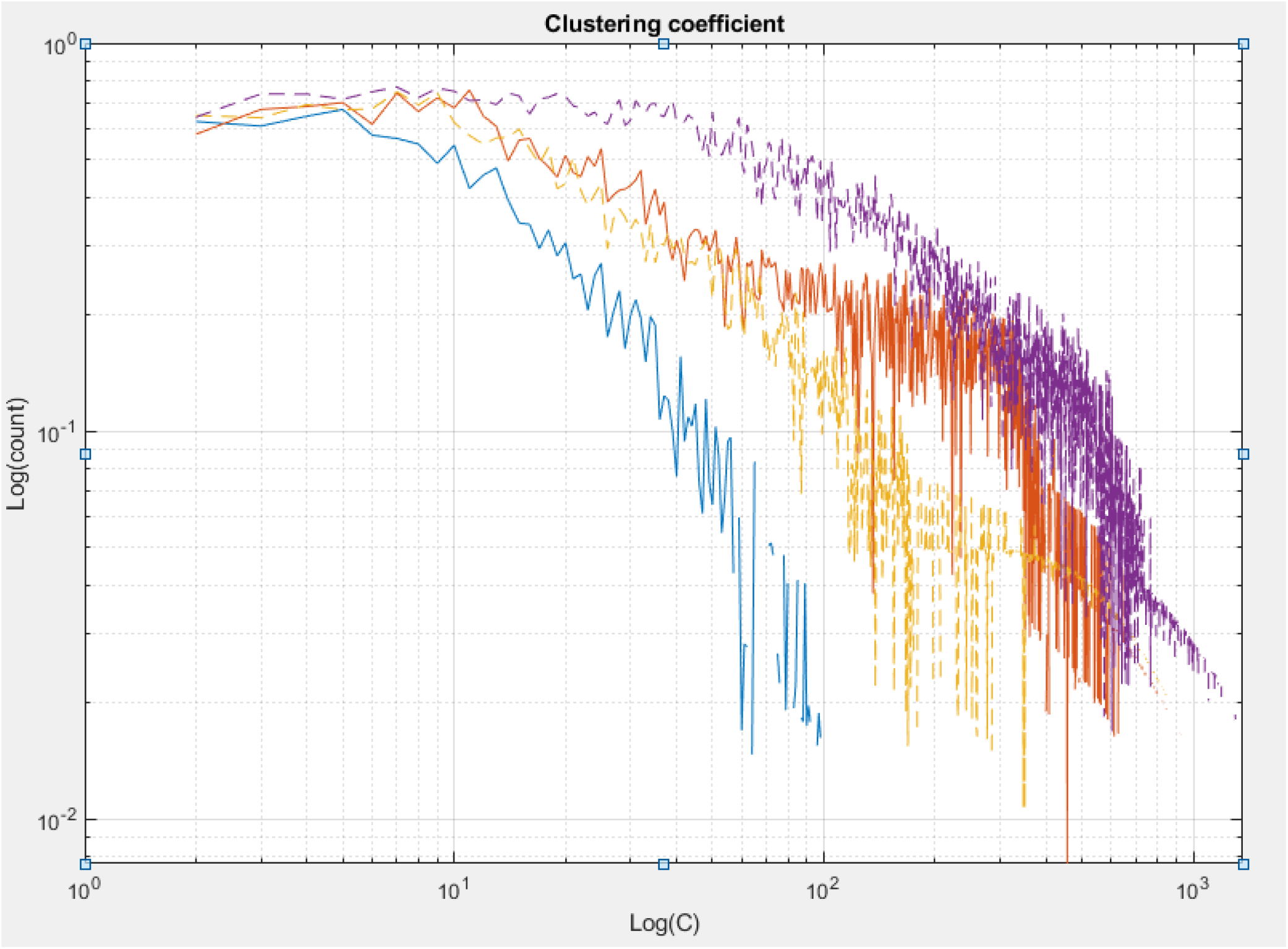
The relation between count and C is revealed in this full-log plot for HS, LS, HW, and LW states. Each state contains 20 samples. Hw and Hs are represented by blue and orange lines, while Lw and Ls are yellow and purple. C4.5 can be used to analyze the clustering distribution to make a new decision tree.

One common method in RS-fMRI (Resting state-fMRI) is Seed-based Correlation Analysis (SCA). SHR and WKY have been studied ^29^. In general, SCA requires the choice of a ROI (Region Of Interest) to be a priori assumption, and needs to average over seed regions before calculating connectivity. In contrast, we forgo this step to avoid any subjective and abnormal seed from affecting the outcome. Instead, we upgrade and simplify the graph theory by numerical representations. Whether this approach is applicable to all states of mental disorder and has operating restrictions are important questions to answer in the future. In this work we have managed to establish that the power-law distribution carries enough information to deal with the trial subjects in this case. To locate the characteristics of any mental state, one only needs to use chi2 algorithm to pick out important degrees filtered from the power-law distribution.

In the past, researchers found that brains have two large opposing systems in the resting state. One is the DMN (Default mode network), while the other is composed of attentional or task-based systems^30^. This motivates us to check whether double power laws may turn out to describe the degree distribution better than the usual simple power law. In other words, can it be that each of these two systems contributes independently and gives rise to two different exponents. In Table 5, we show the outcome of four states and human resting-state by AIC (Akaike information criterion)^31^. AIC is a statistical method to distinguish the best fitting function among multiple candidates. Basically it balances the principles of accuracy (i.e., minimum loss of information) and frugality, as shown in Table 5.

**Table 5.**
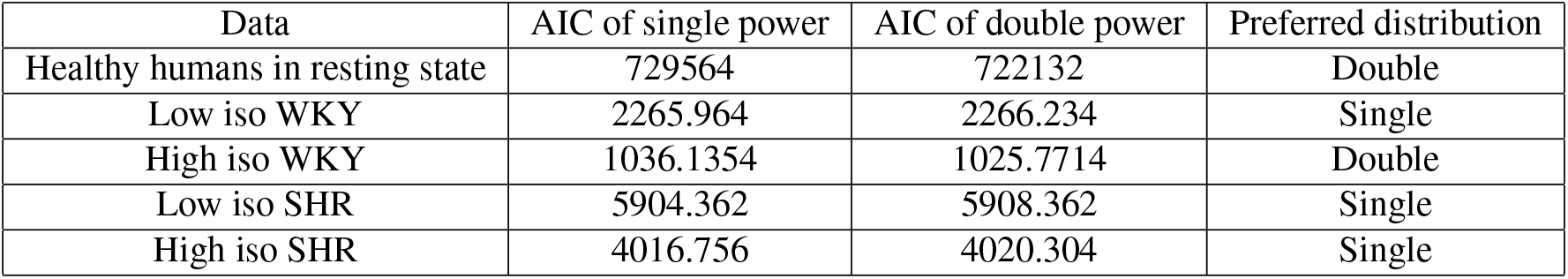
AIC values are used to select the preferred probability distribution for five mental states.

| Data | AIC of single power | AIC of double power | Preferred distribution |
| --- | --- | --- | --- |
| Healthy humans in resting state | 729564 | 722132 | Double |
| Low iso WKY | 2265.964 | 2266.234 | Single |
| High iso WKY | 1036.1354 | 1025.7714 | Double |
| Low iso SHR | 5904.362 | 5908.362 | Single |
| High iso SHR | 4016.756 | 4020.304 | Single |

Table 5 includes five different mental states: (1) Healthy humans in resting-state for which double power laws fit better. (2) Single power law wins out by a small margin for low iso rats, It is worth noting that this result should be treated with cautions because LW and LS rats are hard to remain still during fMRI scanning. (3) High iso WKY rats also favor the double powers. (4, 5) Single power law is a better fit for SHR rats. Recent studies found that SHR (ADHD) children usually exhibit abnormal DMN network^32^. It has also been reported that mental disorder such as Alzheimer^33,34^, depression^35^, schizophrenia^36^, and ASD^37^ can render DMN abnormal. It remains a pressing task to clarify whether the transition of double powers to single power correlates with abnormal DMN. In summary, power-law distribution can not only reflect the mental condition of our samples, but also reveal detail information about their network properties.

It is desirable to have more samples to optimize our use of Decision Tree to select power law from MRI data of mentally disordered rats. Although our results have demonstrated that the power-law distribution can be analyzed by Decision Tree to classify dosage of IOS and SHR vs. WKY, to vindicate its versatility more propositions are needed, e.g., depression, hypertension, or transient ischemic attack. We have two ideas to improve current understandings of the dynamics of brain: first, establish a relationship between observers (i.e., Decision Tree) and the objects being observed (i.e., power law from different states.). Once this relationship is available, it may function as a starting point to reveal possible connections among different observed objects.

## Materials and Methods

### Process

Step 1: Using Pearson correlation to transform 4D fMRI to power law distribution. Step 2: Using chi2 algorithm to select important features from degrees. Step 3: Using the outcome of step 2 as an input to c4.5 to do training and testing. Step 4: Selecting the better one with accuracy over 0.9.

### Animals: Rat

All the experimental animals were admitted by the National Tsing Hua University Institutional Animal Care and Use Committee and complied with experimental guidelines. The important information of rat is listed in Table 6.

**Table 6.** Information of rat

| state | number | age (week) | isoflurane |
| --- | --- | --- | --- |
| Low iso WKY | 20 | 4 – 8 | $\leq 2.0\%$ |
| High iso WKY | 20 | 4 – 8 | $\geq 2.0\%$ |
| Low iso SHR | 20 | 4 – 10 | $\leq 2.0\%$ |
| High iso SHR | 20 | 5 – 10 | $\geq 2.0\%$ |

### Animals: Human

We used six normal human subjects to test single and double power law from Teng-Yi Huang’s Lab. The ADHD database are available to download through the ADHD-200 Consortium[a, b]. All resting-state fMRI scans were performed in New York University Child Study Center [b]. Our retrospective study using the ADHD-200 database was approved by the institutional review boards of National Taiwan University that confirmed both our methods and experimental protocols were in accordance with its guidelines and regulations. Informed consent was obtained from all participants or, if participants are under 18, from a parent and/or legal guadian. [a] The ADHD4-200 Consortium, Front Syst Neurosci. 2012; 6:62 [b] http://fcon_1000.projects.nitrc.org/indi/adhd200/index.html [c] Growing Together and Growing Apart: Regional and Sex Differences in the Lifespan Developmental Trajectories of Functional Homotopy, https://www.jneurosci.org/content/30/45/15034.long

### Magnetic resonance imaging

Our raw data come from the same source in Ref.29, in this section, copyright is owing to the author who wrote ‘‘Magnetic resonance imaging” in the Ref.15. We scanned all animals with 7-Tesla Bruker Clinscan, which had a volume coil for signal excitation and a brain surface coil for signal receiving. The anesthesia process is operated by 1.4–1.5 % isoflurane mixed with O2 at flow rate of 1 L/minute. We monitored all rats, made sure the respiratory rate in the range of 65-75 breaths/min while the scanning period, and body temperature maintained at 37 °C by a temperature-controlled water circulation machine. During the rs-fMRI experiments, we used gradient echo echo-planar-imaging (EPI) getting the 300 consecutive volumes with 11 coronal slices. The EPI specification is TE/TR = 20 ms/1000 ms, matrix size is 64 × 64, FOV= 30 × 30*mm*^2^ and slice thickness = 1 mm. We get the anatomical images by turbo-spin-echo (TSE) with scanning parameters of TE/TR= 14/4000, matrix size = 256 × 256, FOV= 30 × 30*mm*^2^, slice thickness = 1mm, number of average = 2. To inspect the result of deep anesthesia, we applied 2.5 2.7% isoflurane mixed with O2, and monitored respiratory rate in the range of 40–45 breaths/min during the whole scanning period.

### Data processing for distribution of degree

Here we analyze our raw data from fMRI Grayscale image. Afterward, we transform them to scalar value matrix by MATLAB 2015a and 2018 version. This matrix is four dimensional, 64 × 64 × 11 × 525 where the first three components denote spatial position, while the last component refers to the time section in the scanning. Four dimensions render the matrix hard to manipulate, and it costs a lot of computer time. One can use GPU computing to disassemble it to two-dimensional form (45056 × 525). After using Eq.1 to calculate all voxels of degree in whole brain, we can get the distribution of degree for any sample. We calculate all 80 rats (each state has 20 samples) in order to obtain the degree distribution form.

### Chi2 algorithm

The feature selection on this study stems from chi2 algorithm^38^ which is designed to discretize numeric attributes based on the *X*^2^ statistic, and consists of two phases. In the first phase, it begins with a high significance level (sigLevel), Phase 1 is, as a matter of fact, a generalized version of ChiMerge of Kerber. Phase 2 is a ner process of Phase 1. Starting with sigLevel0 determined in Phase 1, each attribute i is associated with a sigLevel[i], and takes turns for merging. Consistency checking is conducted after each attribute’s merging. At the end of Phase 2, if an attribute is merged to only one value, it simply means that this attribute is not relevant in representing the original data set. As a result, when discretization ends, feature selection is accomplished.

### Data processing for C4.5 decision tree

In this study, we propose a set of new algorithms to enhance the Identifying effectiveness of SHR and WKY. The proposed classifier algorithms are a combination of chi2 algorithm and C4.5 decision tree (C4.5), the chi2 algorithm evaluates the worth of a subset of attributes and C4.5 speculate the mental disorder. The chi2 algorithm is commonly used for testing relationships between categorical variables. The calculation of the chi2 algorithm is follows 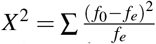, where *f*_0_ = the observed frequency (the observed counts in the cells) and *f_e_* = he expected frequency if NO relationship existed between the variables.The decision tree algorithm is well known for its robustness and learning efficiency with a learning time complexity of *O*(*nlog_2_n*)^39^, C4.5 has been listed in the top 10 algorithms in data mining ^40^. It is a popular statistical classifier developed by Ross Quinlan in 1993. Basically, C4.5 is an extension of Quinlan’s earlier ID3 algorithm. In C4.5 the Information Gain split criterion is replaced by an Information Gain Ratio criterion which penalizes variables with many states. C4.5 can be used to generate a decision tree for classification. The learning algorithm applies a divide-and-conquer strategy ^41^ to construct the tree. The sets of instances are accompanied by a set of genes (attributes). This classifier has additional features, such as handling missing values, categorizing continuous attributes, pruning decision trees, deriving rules, endotestae Information gain (S, A) of a feature A relative to a collection of examples S, is defined as 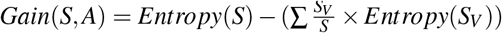, where Values (A) is the set of all possible values for attribute A, and Sv is the subset of S for which feature A has value v (i.e *S_V_* = {*s* ∈ *S* | *A*(*s*) = *v*}), Note the first term in the equation for Gain is just the entropy of the original collection S and the second term is the expected value of the entropy after S is partitioned using feature A. The expected entropy described by the second term is the direct sum of the entropy of each subset Sv, weighed by the fraction of samples 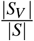 that belong to *S_V_*, Gain (S, A) is therefore the expected reduction in entropy caused by knowing the value of feature A. The Entropy is given by 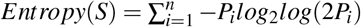.

### Data processing for testing single and double power law

We choose the same method and procedures described in Ref.41. Details can be found in Sec.IV^42^.

### Data processing for calculating L

When we calculated the degree for any voxels, the path length (L) between two voxels is defined as the minimum number of links necessary to connect each other. Also, we collect L from data of six rats for each state at the same time. Afterward, MATLAB 2018a is employed to find the max path and average L.

### Data processing for C and structure network patterns

First, we print out all connection information between two voxles for any sample in txt format. Then, insert the data in MATLAB 2018a to obtain the functional brain network pattern. If interested at calculating the clustering coefficient for any voxel linked, you have to get information of degree and the number of links connecting the neighbors. Finally, the average C can be determined from this equation 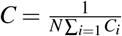 where *N* is the number of voxels and *i* the voxel number.

## Acknowledgements

We acknowledge funding from the Ministry of Science and Technology in Taiwan under grant no 105-2112-M-007-008-MY3 and 108-2112-M007-011-MY3. We also thank Prof. Chung-Chuan Lo for helpful advices, Prof. Che-Rung Lee and Quey-Liang Kao for the assess to their GPU workstation, and Kun-I Chao and Li-Jie Chen for helping us process images of fMRI.

## Author contributions statement

Hosting Plan: Ping-Jui Tsai and Kun-Huang Chen.

Coding program: Ping-Jui Tsai.

Image processing: Ping-Jui Tsai.

Data processing: Ping-Jui Tsai.

Statistical Analysis: Kun-Huang Chen (for chi2 algorithm, C4.5 decision tree, t-test, and ROC), Ping-Jui Tsai (for AIC and power law).

Academic guidance: Fu-Nien Wang, Tzay-Ming Hong and Teng-Yi Huang.

Writing: Ping-Jui Tsai and Tzay-Ming Hong (Abstract, Introduction, Result, Discussion), Ping-Jui Tsai and Kun-Huang Chen (Materials and Methods).

## Competing interests

**The authors declare no competing interests**.

All authors reviewed the manuscript.

